# A non-parametric method to compute protein–protein and protein–ligands interfaces. Application to HIV-2 protease–inhibitors complexes

**DOI:** 10.1101/498923

**Authors:** Pierre Laville, Juliette Martin, Guillaume Launay, Leslie Regad, Anne-Claude Camproux, Sjoerd de Vries, Michel Petitjean

**Affiliations:** Université Paris Diderot, MTi, INSERM UMR-S 973, Paris, France; Université Lyon 1, IBCP, MMSB, CNRS UMR 5086, Lyon, France; Université Paris Diderot, E-pôle de génoinformatique, Institut Jacques Monod, CNRS UMR 7592, Paris, France

## Abstract

**Motivation:** The accurate description of interfaces is needed to identify which residues interact with another molecule or macromolecule. In addition, a data structure is required to compare interfaces within or between families of protein–protein or protein–ligands complexes. In order to avoid many unwanted comparisons, we looked for a parameter free computation of interfaces. This need appeared at the occasion of bioinformatics studies by our research team focusing on HIV-2 protease (PR2) resistance to its inhibitors.

**Results:** We designed the PPIC software (Protein Protein Interface Computation). It offers three methods of computation of interfaces: (1) our original parameter free method, (2) the Voronoi tessellation approach, and (3) the cutoff distance method. For the latter, we suggest on the basis of 1050 dimers protein–protein interfaces that the optimal cutoff distance is 3.7 Å, or 3.6 Å for a set of 18 PR2–ligand interfaces. We found at most 17 contact residues with PR2 ligands.

**Availability:** Free binaries and documentation are available through a software repository located at http://petitjeanmichel.free.fr/itoweb.petitjean.freeware.html

## 1 Introduction

As noticed by Vangone and Bonvin (2015), the number of connections between each pair of proteins is a strong predictor of how tightly the proteins connect to each other. According to Janin et al. (2008), de Vries and Bonvin (2008) and Dequeker et al. (2017), there are three main methods to compute interfaces between protein chains:

1. The cutoff method (implemented in PPIC).
2. The loss of ASA (accessible surface area) upon binding.
3. The Voronoi tessellation method (implemented in PPIC).

The cutoff method assumes that two residues are in contact when they have a pair of heavy atoms separated by a distance smaller than the cutoff distance. This cutoff distance may be set to 4.5 Å between non H atoms (Gao and Skolnick, 2010; Esque et al., 2011; Triki et al., 2018), to 5 Å from side-chain centers of mass (Viloria et al., 2017), or to 8 Å between C_α_ atoms (Esque et al., 2011). The choice of the cutoff can significantly influence the result (de Vries and Bonvin, 2008). This method critically relies on the choice of atomic radii, which strongly depend on how they are defined (Bondi, 1964; Allinger et al., 1994). The values recommended in the literature (Gavezzotti, 1983) induce the existence of intersections of 6 spheres or more. These intersections above 3 or 4 spheres are ignored by a number of published methods, inducing in turn large errors in surface calculations (Petitjean, 1994, 2013).

In its original variant, the Voronoi tessellation method is parameter-free (Cazals et al., 2006; Bouvier et al., 2009). It was implemented in PROVAT (Gore et al., 2005). The full mathematical presentation of Voronoi diagrams and their calculation algorithms are out of the scope of this paper (see Edelsbrunner, 1987). We may just say that each atom lies inside a convex polyhedral cell having its polygonal faces located at mid distance from its neighbouring atoms. Thus, two atoms are neighbours if their Voronoi cells share a common face. Extensions taking in account atomic spheres were proposed (Esque et al., 2011; Mahdavi et al., 2012), but these latter are no more parameter-free.

## 2 Methods

Our original method, which is implemented in PPIC, is a variant of the one of Cerisier et al. (2017). The input is a complex with two partners (molecule or macromolecule), A and B. The algorithm has no parameter. It has two steps:

1. Generate the interface in A as the non-redundant set of all nearest neighbours in A of the atoms of B.
2. Generate the interface in B as the non-redundant set of all nearest neighbours in B of the atoms of A.

The interface has two parts, one in A and one in B. Each one is a subset of the interface that would be output by the Voronoi tessellation method (proof: see Eppstein et al., 1997). However our method is much simpler that the latter, and it does not generate neighbours at long distances in the interface, while large Voronoi cells can induce such long distances. Furthermore, looking at interacting atom pairs among the nearest neighbours makes more sense that looking at interacting atom pairs at farther distances that the nearest neighbours.

## 3 Results and discussion

We considered the dimers set of Martin et al. (2008), containing 1050 protein homo- and heterodimers. We compared the 1050 couples of interfaces with the ones obtained with the cutoff distance method, for a cutoff ranging from 2 to 6 Å. For the full database, computing these interfaces took about 288 min on a workstation Ubuntu 18.04 (16 Intel^®^ Xeon^®^ CPUs, 3 GHz). This computing time was roughly the same for the three methods implemented in PPIC. The subsets of atoms defining the interfaces were compared with the symmetric difference metric (Deza and Deza, 2009). Similarly, we compared the 36 couples of interfaces for the set of 18 PR2–ligand interfaces from Triki et al. (2018). Figure 1 shows that the closest results for the two methods were observed at a cutoff of 3.7 Å for the dimers set, and at 3.6 Å for the PR2 complexes set. This is a remarkable agreement since these two data sets are of different nature. It suggests an optimal cutoff value of 3.6–3.7 Å between heavy atoms. This optimal value is in agreement with the statistical analysis of McDonald and Thornton (1994): the distance donor–acceptor should be less than 3.9 Å and the distance hydrogen–acceptor should be less than 2.5 Å. So, assuming a covalent H bond length of 1.1 Å, the distance donor–acceptor should be in fact at most 2.5 + 1.1 = 3.6 Å, which is our suggested value. For the Voronoi tessellation method, the closest results were observed at the respective cutoffs of 5.1 Å and 4.6 Å. In the case of distances between heavy atoms, these cutoff values are too large, as predicted in section 2.

For the PR2 complexes, we found that at most 17 residues are in contact with the ligands: Arg 8, Leu 23, Asp 25, Gly 27, Ala 28, Asp 29, Asp 30, Ile 32, Ile 46, Val 47, Gly 48, Gly 49, Ile 50, Phe 53, Pro 81, Ile 82 and Ile 84. All were found in the binding pocket computed by the consensus method of Triki et al. (2018) based on a 4.5 Å cutoff distance. This confirms the pertinence of our parameter-free method.

**Figure 1:**
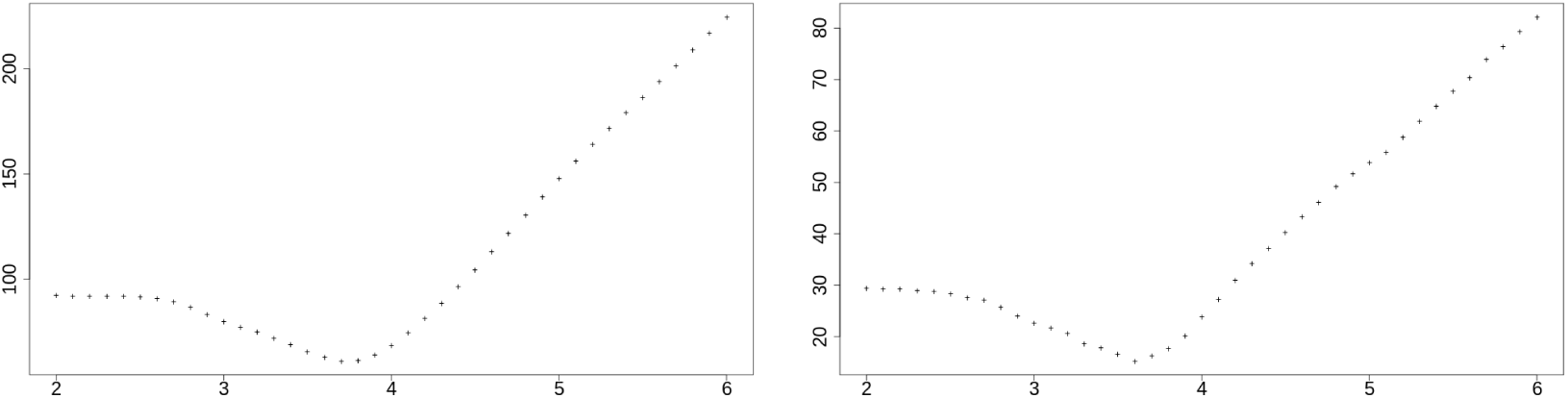
Mean symmetric difference distances in function of the cutoff value, for the 1050 dimers data set (on the left) and for the 18 PR2 complexes (on the right).

